# Genotyping of Russian isolates of fungal pathogen *Trichophyton rubrum*, based on simple sequence repeat and single nucleotide polymorphism

**DOI:** 10.1101/2020.03.06.980839

**Authors:** Ivan M. Pchelin, Yuri V. Mochalov, Daniil V. Azarov, Sofya A. Romanyuk, Galina A. Chilina, Irina V. Vybornova, Tatiyana V. Bogdanova, Vasily V. Zlatogursky, Svetlana V. Apalko, Natalia V. Vasilyeva, Anastasia E. Taraskina

## Abstract

**Background:** The *Trichophyton rubrum* species group consists of prevalent causative agents of human skin, nail and hair infections, including *T. rubrum sensu stricto* and *T. violaceum*, as well as other less well established or debatable taxa like *T. soudanense*, *T. kuryangei* and *T. megninii*. Our previous study provided limited evidence in favour of the existence of two genetic lineages in the Russian *T. rubrum sensu stricto* population.

**Objectives:** We aimed to study the genetic structure of the Russian population of *T. rubrum*, and to identify factors shaping this structure.

**Methods:** We analysed the polymorphism of 12 simple sequence repeat (SSR, or microsatellite) markers and single-nucleotide polymorphism in the TERG_02941 protein-coding gene in 70 *T. rubrum* isolates and performed a phylogenomic reconstruction.

**Results:** All three types of data provided conclusive evidence that the population consists of two genetic lineages. Clustering, performed by means of microsatellite length polymorphism analysis, was strongly dependent on the number of nucleotide repeats in the 5’-area of the fructose-1,6-bisphosphate aldolase gene. Analysis of molecular variance (AMOVA) on the basis of SSR typing data indicated that 22–48% of the variability was among groups within *T. rubrum*. There was no clear connection of population structure with types of infection, places of geographic origin, aldolase gene expression or urease activity.

**Conclusion:** Our results suggest that the Russian population of *T. rubrum* consists of two cosmopolitan genetic lineages.

## Introduction

*Trichophyton rubrum* is an ascomycete from the order Onygenales. It is prevalent fungus of medical importance in most countries due to its major contribution to the etiology of toenail onychomycosis and foot skin mycosis.^1^ *Trichophyton rubrum sensu stricto* is a part of species group, which also includes *T. violaceum* and *T. soudanense* along with several forms, potentially deserving species status. All of these can be differentiated by ribosomal DNA internal transcribed spacer (ITS) region sequences.^2–6^ Statistical analysis of genetic polymorphism in simple sequence repeats (SSR, or microsatellites) revealed clonal mode of reproduction in the species.^7–8^ It is expectedly accompanied by low genetic polymorphism.^9–11^ Several studies of mating type locus idiomorphs distribution in *T. rubrum* population found a single mating type locus variant, out of two, which are necessary for mating.^12–14^ A certain proportion of *T. rubrum* isolates, belonging to the so-called “raubitschekii” morphotype, possess urease activity. This trait is believed to be ancestral,^15^ but has unclear value for taxonomy.

To the best of our knowledge, by the moment there are two papers, reporting bipartite population structure in *T. rubrum sensu stricto*. In a previous study, we reported a concordance between ribosomal non-transcribed spacer region (NTS) genotypes and nonsynonymous single nucleotide polymorphisms (SNPs) in two protein-coding loci, TERG_02941 and TERG_03298 in *T. rubrum* isolates from a city in North-Western Russia.^16^ The two genetic lines were named ST1 and ST2. Also, Zheng et al.^17^ studied a sample of *T. rubrum* isolates, covering major part of China, by whole genome sequencing (WGS). A number of other studies obtained *T. rubrum* WGS data from wide variety of geographic locations, covering Europe, Asia and North America.^4,14,18,19^ Clear understanding of the potential of low throughput methods to uncover genetic population structure can be achieved when their results are cross-validated and compared with whole genome sequence-based trees. However, there have been no studies, bringing together data on SNP, SSR, WGS and NTS polymorphism in *T. rubrum*.

Our study aimed to verify the presence of previously shown genetic structure in Russian population of *T. rubrum sensu stricto*, to find whether this structure is linked to particular characteristics and, finally, to identify factors, which shape the structure.

## Materials and Methods

### Fungal isolates

Clinical isolates of *T. rubrum* have been obtained in the clinic of Kashkin Research Institute of Medical Mycology, Saint Petersburg (n = 44) and in Sverdlovsk Regional Dermatovenereologic Dispensary, Yekaterinburg (n = 23) from October 2015 to January 2018 (Supplement S1) and subcultured on Sabouraud agar medium, with the addition of dextrose to a final concentration of 2%. We also used two reference strains RCPF 1280 and RCPF 1408 and the type strain CBS 392.58 up to a total of 70 strains in the sample. This sample shared 19 isolates with our earlier study.^16^ Major part of strains originated from toenail onychomycosis and tinea pedis cases (87%), 6% were from tinea corporis, 4% from tinea cruris and 3% from fingernail onychomycosis. The following clinical *T. rubrum* isolates were deposited in the Russian Collection of Pathogenic Fungi (RCPF): D15P33, RCPF 1864; D15P44, RCPF 1865; D15P50, RCPF 1866; D15P62, RCPF 1867; D15P71, RCPF 1868; Y1013, RCPF 1869; D15P139, RCPF 1870. The urease activity was tested twice in Christensen’s urea broth, supplemented with chloramphenicol to a final concentration of 40 μg/ml.

### Species identification

Species identification was performed by phenotypic methods and confirmed by DNA sequencing of ITS region. To extract genomic DNA, we used GeneJET Plant Genomic DNA Purification Mini Kit (Thermo Fisher Scientific, Lithuania). Aerial mycelium and fungal spores were scraped by pipette tips, placed into microtubes containing Lysis Buffer A, beaten with glass beads in Minilys tissue homogenizer (Bertin Technologies, France), supplemented with Lysis Buffer B and RNase A and further treated according to the kit manufacturer’s protocol. Amplification of ITS region with ITS5 and ITS4 primers^20^ (Table 1) was performed in 50-μl volumes, using standard kit for PCR diagnostics (Syntol, Russia). Reaction mixture consisted of 100 mM Tris/HCl (pH 9.0), 250 mM KCl, 10 mM Tween-20, 2.5 mM MgCl_2_, 250 μM each dNTP and contained 5 pmol of primers, 1 U Taq polymerase, and 2.5 μl of template DNA. The thermal cycler C1000 Touch (Bio-Rad, USA) was programmed as follows: initial denaturation step for 5 min at 95 °C, 34 cycles of amplification: denaturation for 30 s at 95 °C, annealing 30 s at 56 °C, extension for 1 min at 72 °C, and a final extension step for 10 min at 72 °C. The products of PCRs were purified by isopropanol precipitation in the presence of glycogen. PCR products were sequenced on both strands using BigDye Terminator v3.1 Cycle Sequencing Kit (Thermo Fisher Scientific, USA) on a 3500 Genetic Analyzer (Thermo Fisher Scientific, USA).

**Table 1.**
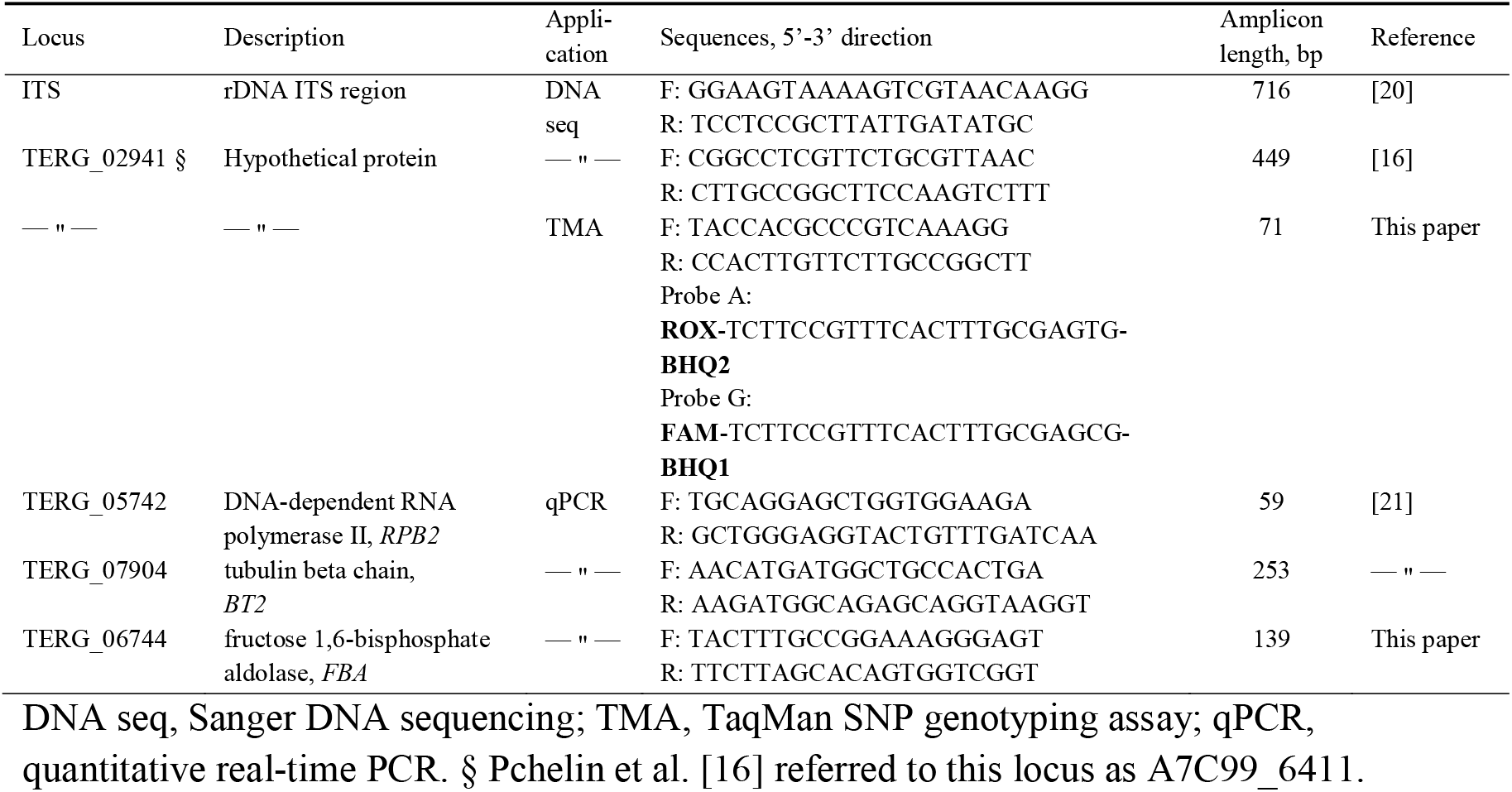
Primers, used in the study.

### Antifungal susceptibility testing

Antifungal susceptibility testing was performed according to EUCAST E.Def 9.3.1 document,^22^ in RPMI-1640 medium with the addition of glucose to a final concentration of 2%. We tested the activity of terbinafine, fluconazole and itraconazole against six *T. rubrum* isolates with newly obtained WGS sequences. Terbinafine was tested at concentration range 0.007-4 *μ*g/ml, fluconazole, itraconazole and voriconazole were tested at 0.015-8 *μ*g/ml, posaconazole was tested at 0.008-4 *μ*g/ml.

### Molecular strain typing

We employed two strain typing approaches. Firstly, we probed the G793A nucleotide substitution in protein-coding locus TERG_02941 (A7C99_6411) by Sanger DNA sequencing. The amplification was carried out under the following conditions: 5 min at 95 °C; 39 cycles of 30 s at 95 °C, 1 min 5 s at 64 °C; and final elongation for 5 min at 72 °C. All other steps were performed as described in the section 2.2 for ITS region sequencing. To optimize the genotyping process, we developed TaqMan SNP genotyping assay with two allele-specific probes (Table 1). Samples were run using a C1000 Touch thermal cycler equipped with a CFX96 real-time PCR detection system (Bio-Rad, USA) according to the following program: initial denaturation step for 3 min at 95 °C; and 39 cycles of 10 s at 95 °C and 30 s at 59 °C. The assay generated fluorescent signal from both probes in every sample, specific signal reached the threshold value with 4-cycle outrunning.

Secondly, we performed microsatellite typing. We used eight microsatellite loci from the papers by Gräser et al.^7^ and Gong et al.^8^ and developed four additional loci (Table 2). To find new polymorphic loci, we searched genome contigs for repeat sequences and prepared nucleotide alignments. In total, 170 loci were screened. Primers for amplification of nine candidate loci were designed with the use of Primer-BLAST tool.^23^ Five loci had uniform lengths in 21-56 isolates and were excluded from further experiments. Amplification of microsatellite loci was performed in 25-μl volumes. The primers used are listed in the Table 2.

**Table 2.**
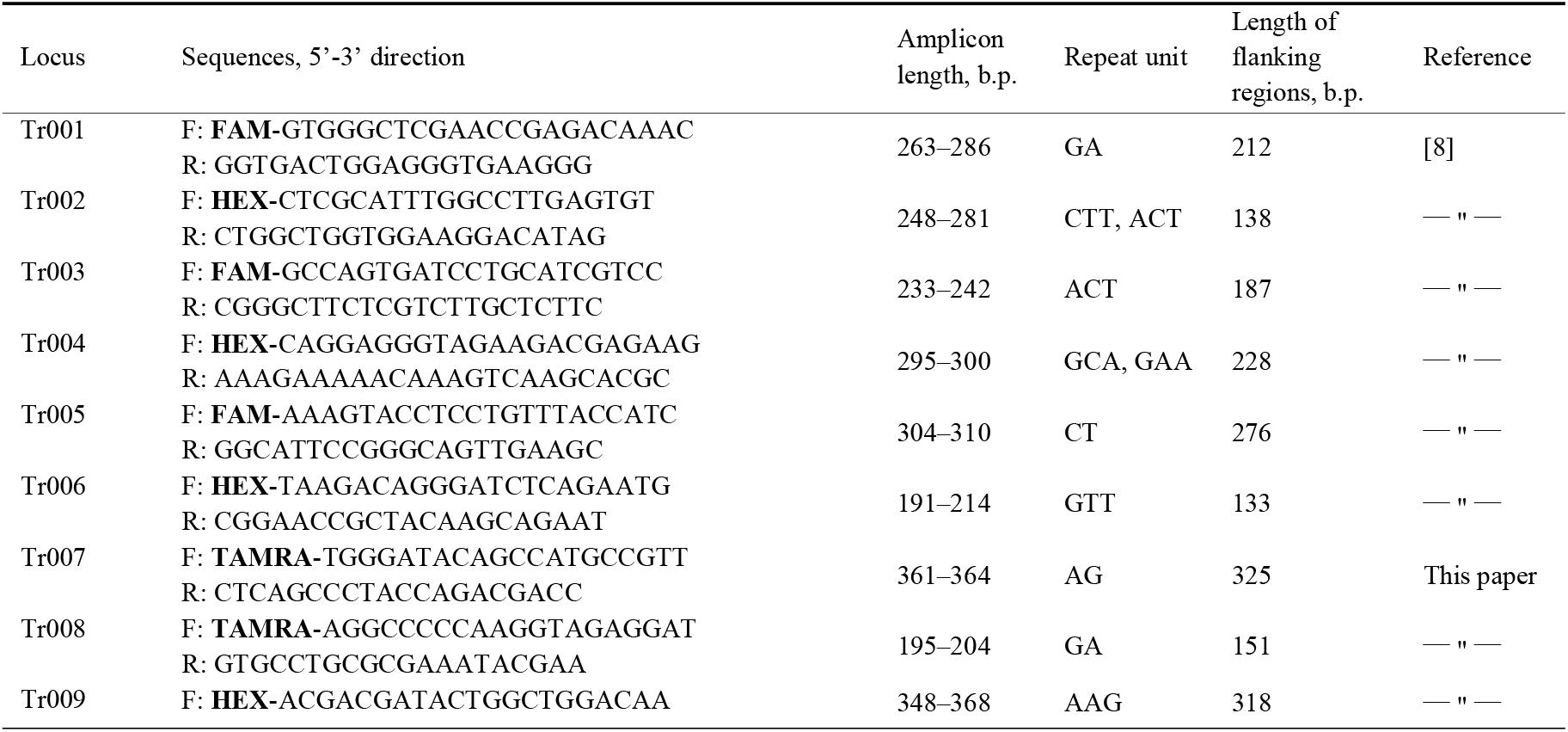
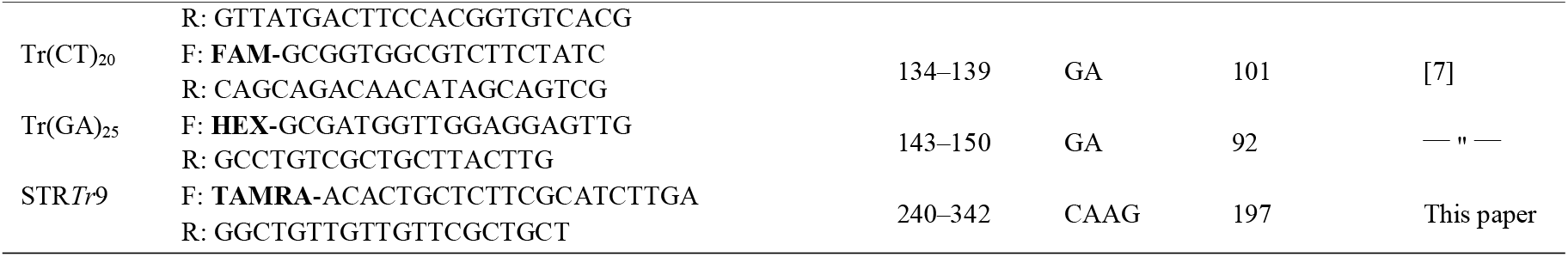
Primers, used for SSR amplification in *Trichophyton rubrum* strains. Fluorescent modifications are given in bold.

The amplification program for microsatellites included initial denaturation at 95 °C for 10 min, followed by 30 cycles of 50 s at 95 °C, 1 min at 58 °C, and 1 min at 72 °C, and a terminal extension step of 72 °C for 10 min. Fluorescent PCR products were subjected to fragment analysis on a 3500 Genetic Analyzer (Thermo Fisher Scientific, USA) in the presence of Red DNA Size Standard (Molecular Cloning Laboratories, USA). The set of fluorescent dyes used in this work appeared to be suboptimal because of the need to check data for issues of spectral overlapping between TAMRA and ROX channels.

### Whole genome sequencing

Genomic DNA of six *T. rubrum* isolates was extracted using GeneJET Plant Genomic DNA Purification Mini Kit (Thermo Fisher Scientific, Lithuania). Preparation of DNA sample library was done with the use of Kapa HTP Library Preparation Kit (Roche, USA), according to the manufacturer’s instructions. Paired end sequencing (250 × 250 cycles) was performed using the MiSeq sequencer (Illumina, USA). The reads obtained were prepared for the analyses as described earlier.^24^ Quality control of the reads was performed using FastQC 0.11.8.^25^ It was followed by trimming low-quality reads and TruSeq2 adapters with Trimmomatic 0.39.^26^ De novo assembly was performed with SPAdes 3.9.0 with the “--careful option”.^27^ Raw reads were submitted to the Sequence Read Archive of National Center for Biotechnology Information (BioProject ID PRJNA552357).

### Phylogenomic reconstruction

We analyzed six original whole genome sequences, as well as the genome of isolate CBS 118892, NCBI BioProject accession ACPH00000000;^18^ isolates CBS 202.88, AOKX00000000; CBS 288.86, AOKU00000000; CBS 289.86, AOKV00000000; CBS 100081, AOKT00000000; D6, AOLB00000000; MR850, AOKR00000000; MR1448, AOKZ00000000; MR1459, AOLC00000000;^14^ isolate CMCC(F)T1i, LHPM00000000;^4^ IGIB-SBL-CI1, JPGR00000000.^19^ The raw read data for eight strains from the project of Zheng et al.^17^ have been downloaded from NCBI SRA database and assembled *de novo* for the analysis. This number included the strains LZ164 (NCBI SRA accession SRX5814844), GZ0580 (SRX5814815), NJ785 (SRX5814822) from one cluster, and the strains LZ155 (SRX5814845), NJ786 (SRX5814823), NJ823 (SRX5814824), DL7506 (SRX5814836) from another cluster, revealed in the original publication. The eighth strain GZ0663 (SRX5814819) from the study of Zheng et al., analyzed here, had unclear cluster affiliation. The sequences of *T. rubrum* morphotype “megninii” CBS 735.88 JHQM00000000, *T. soudanense* CBS 452.61 AOKW00000000 and *T. violaceum* CMCC(F)T3l LHPN00000000 were used as an outgroup. Single-nucleotide polymorphism analysis in core genome was done in Harvest Parsnp 1.2 software.^28^ Parsnp run with *T. rubrum* CBS 118892 assembly GCA_000151425.1 as a reference and with default parameters. Whole-genome phylogenetic tree was reconstructed by Maximum Likelihood method in FastTree2,^29^ as implemented in Harvest suite. Branch support values were calculated by performing Shimodaira-Hasegawa test on three alternate topologies around particular splits, with a total of 1000 resamples.

### Analysis of genetic polymorphism

A matrix of mutation probability-based genetic distances^30^ was calculated in polysat 1.7-3 package for R^31^ using microsatellite polymorphism data. Distances in terms of weighted divergence of microsatellite loci were calculated in Microsoft Excel 2010 according to the formula, modified for haploids from.^32^ The dissimilarity *d_i_* between haplotypes *A* and *B* at particular locus *i* was calculated as absolute value of arithmetic difference between the lengths of PCR products *A_i_* and *B_i_*

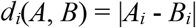

The genetic distance *D* between haplotypes *A* and *B* was based on the sum of dissimilarities *d* between haplotypes at all loci, weighted by dividing this sum by the number of loci taken into analysis, *k*

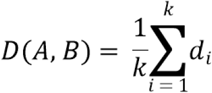

Both distance matrices were further employed in the construction of phylogenetic network in SplitsTree 4.14.2 program.^33^ Neighbor-Joining tree on the basis of Bruvo distances was calculated in Mega 5.2.^34^

The allele association index was calculated from the lengths of microsatellite loci in poppr 2.8.1 package for R,^35^ using the Multilocus Style Permutation method, after deleting one non-informative locus and performing clonal correction. The null hypothesis was the absence of association between the lengths of microsatellite loci. Simpson’s index of genotype diversity within upper and lower bounds for a 95% confidence interval was calculated in the package polysat on the basis of Bruvo genetic distances, in equal groups of 21 isolates for ST1 and ST2 lineages. The analysis of molecular variation (AMOVA) was done on the basis of Bruvo and Rozenfeld distance matrices in GenAlEx 6.5 package^36,37^ with 999 standard permutations.

TERG_03298 genotypes, MAT idiomorphs and TERG_05790 urease gene variants were determined through Blast searches in whole genome sequences.^38^ For mating type determination, TERG_02406 gene coding for α-domain protein was probed by AB759084 sequence.^13^ The absence of HMG protein-coding gene was verified by the searches with *T. simii* HMG gene, GenBank accession number AB605766.^39^

### Quantitative RT-PCR

We hypothesized the influence of number of STR*Tr*9 repeats on *FBA* gene expression under normal conditions. The sample of six *T. rubrum* isolates was grown on Sabouraud agar medium for two weeks at 28 °C. RNA was isolated with the use of GeneJet Plant RNA Purification Mini Kit (Thermo Fisher Scientific, Lithuania). The quantitative RT-PCR experiments were performed in triplicates on two independently grown colonies for each isolate in the sample with the use of OneTube RT-PCR SYBR Kit (Evrogen, Russia). The reactions were carried out on CFX96 Touch Real-Time PCR Detection System (Bio-Rad, USA) with the following cycling conditions: reverse transcription step at 55 °C for 15 min, polymerase activation and reverse transcriptase inactivation step at 95 °C for 1 min, and main amplification stage for 40 cycles, including denaturation at 95 °C for 15 s, annealing at 60 °C for 20 s and elongation at 72 °C for 15 s. Melting curve analysis was performed to ensure that a single PCR product was obtained. Relative gene expression levels were assessed using beta-tubulin and DNA-dependent RNA polymerase II for normalization. The sequences of primers used are given in the Table 1.

## Results

The majority of 69 *T. rubrum* isolates from Russia and the type strain CBS 392.58 had the same ribosomal ITS region sequence, identical to KT285224. There were two exceptions, the isolate D15P139 had 247insC mutation, and another isolate had additional TA element in the microsatellite near the 3’-end of the sequence. Our preliminary study showed discrepancies between the results of strain typing by TERG_02941 locus polymorphism and the microsatellite typing assay of Gong et al.,^8^ based on eight loci.^40^ We hypothesised that this discrepancy was due to insufficient informativeness of the assay and used four additional polymorphic loci, up to a total of 12. The stability of microsatellite loci was assessed in 20 isolates for the time span of 1.5 years. Observed differences of less than one repeat length were considered insignificant, since those differences could be associated with the rounding of measurements, fluctuating near the half-integers. At the endpoint, mutations were observed in five loci. The most extensive changes were noticed in Tr(CT)_20_ locus, with 30% of isolates having mutations. The most pronounced indels were in Tr006 and Tr008 loci, covering up to 3 repeat lengths.

Our first approach for calculating microsatellite polymorphism-based genetic distances employed the probability of mutation events, described with the use of stepwise mutation model.^30^ The split network, inferred using mutation probability-based distances, was divided into three major partitions. One partition contained TERG_02941 793A isolates and another two partitions contained TERG_02941 793G isolates (Fig. 1A). The second approach was based on the comparison of the lengths of microsatellite loci.^32^ The resulting phylogenetic network also had pronounced division into TERG_02941 793A and 793G parts (Fig. 1B). In Rozenfeld distances-based network, further subdivision of TERG_02941 793G partition was less evident, but still discernable. From 22 TERG_02941 793G isolates, 18-21 were located in the same subgroups in both networks.

**Figure 1.**
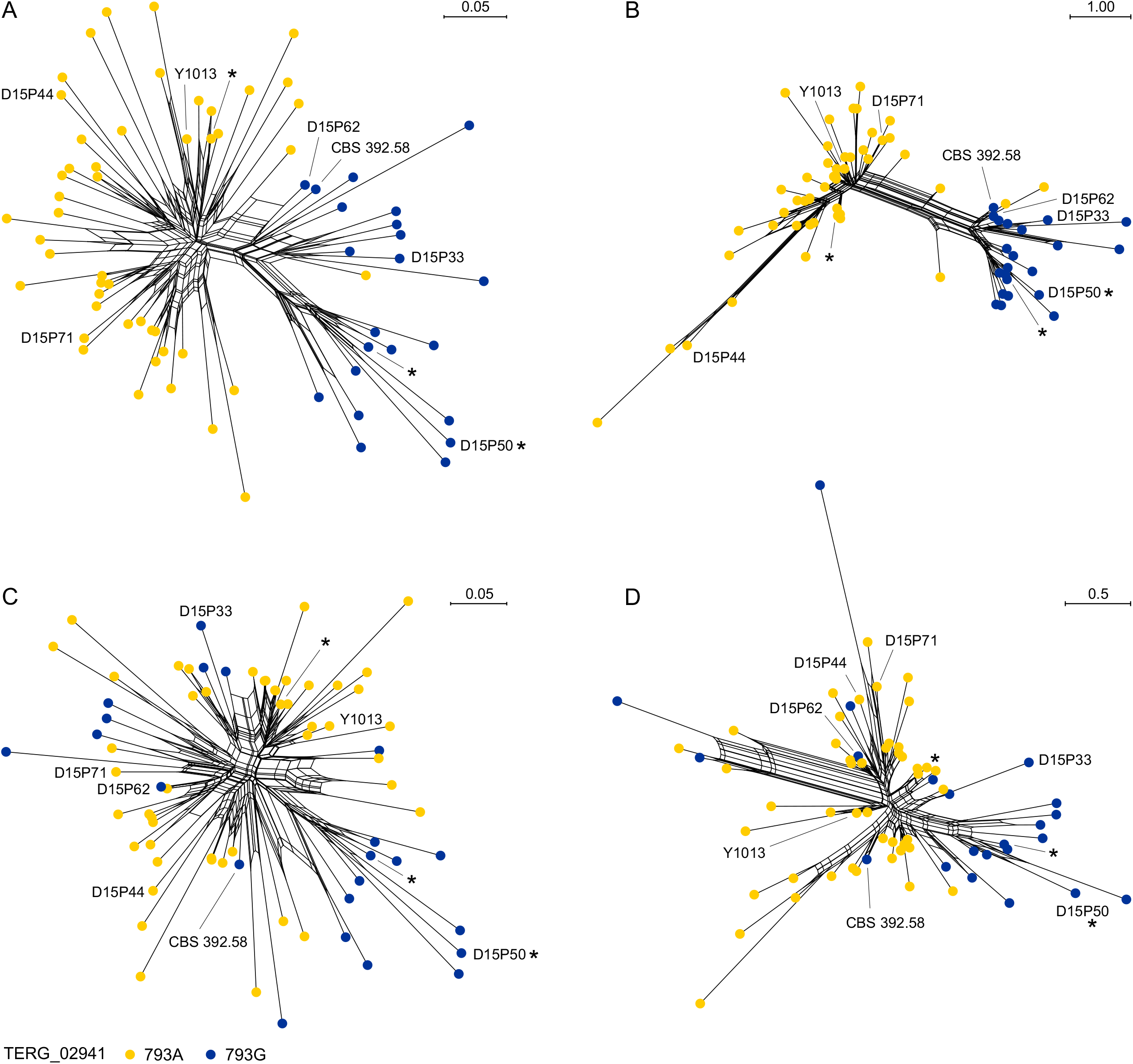
Phylogenetic networks, illustrating the structure of Russian population of *Trichophyton rubrum*, as revealed by simple sequence repeat (SSR) and single nucleotide polymorphism (SNP) genotyping. Isolates with studied whole genome sequences and the type strain CBS 392.58 are indicated. Urease-positive isolates are marked by asterisks. A. Phylogenetic network, calculated from a matrix of mutation probability-based genetic distances (Bruvo), 12 SSR loci. B. Phylogenetic network, calculated from a matrix of distances, based on the lengths of 12 SSR loci (Rozenfeld). The networks C and D were calculated without data on STR*Tr*9 locus polymorphism. C. Phylogenetic network, calculated from a matrix of Bruvo distances, 11 loci. D. Phylogenetic network, based on Rozenfeld genetic distances, 11 microsatellite loci. The removal of STR*Tr*9 locus from the analysis had major influence on the topologies of phylogenetic networks.

For the two genetic lines, the average lengths of PCR products of Tr001, Tr002, Tr003, Tr004, Tr005, Tr007, Tr008, Tr009, Tr(CT)_20_ and Tr(GA)_25_ loci differed by the length of less than one repeat unit. Average Tr006 lengths differed by a single unit length. For STR*Tr*9 locus, this difference was 7.5 unit lengths (Fig. 2A). This difference should be obviously explained by the pressure of natural selection. Since the analysis of intraspecific divergence in most cases is carried out using neutral loci,^41^ we performed calculation of genetic distances without STR*Tr*9 length data. This broke clear grouping of isolates in both phylogenetic networks. However, a certain tendency to clustering of TERG_02941 793G isolates persisted (Figs. 1C, 1D).

**Figure 2.**
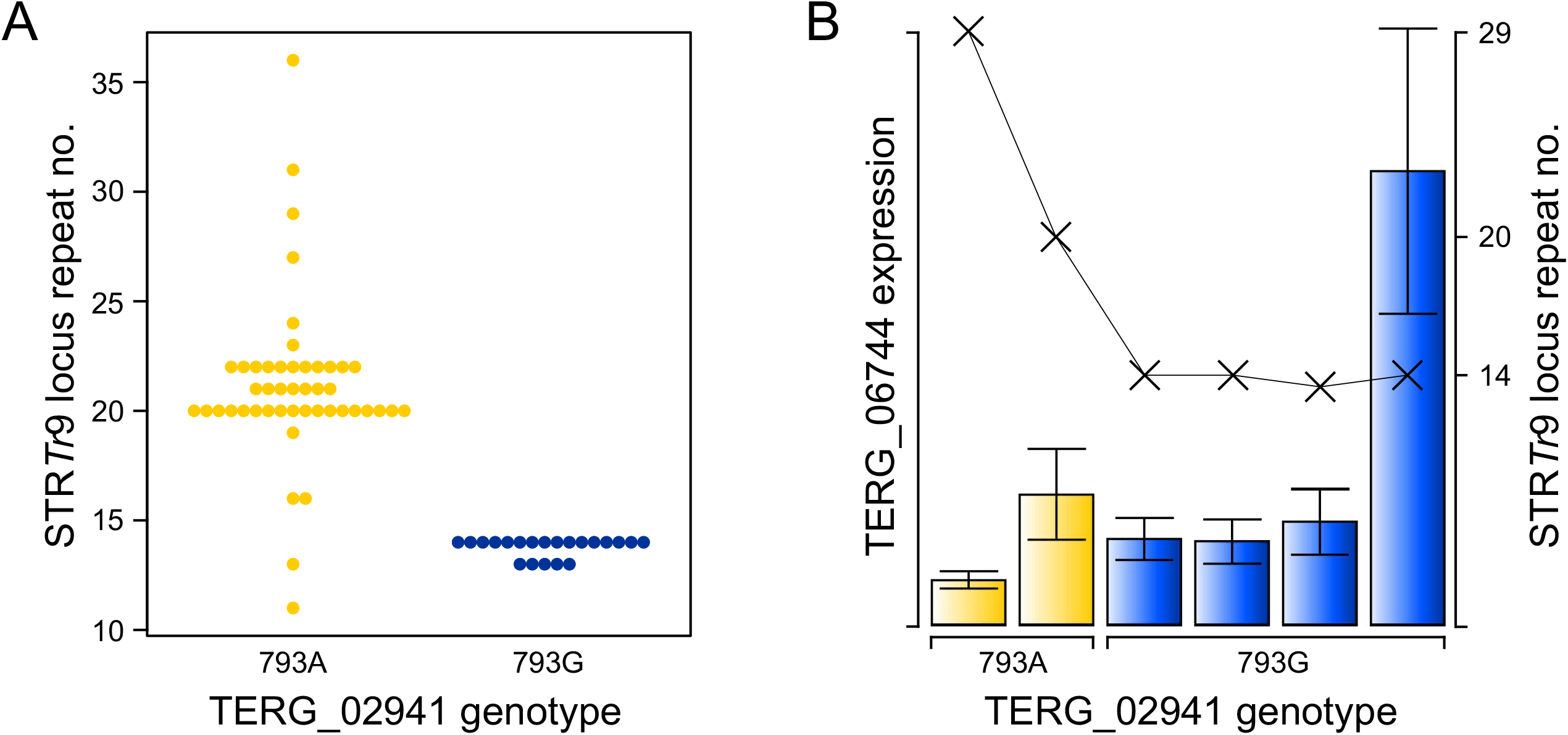
The number of repeats in STR*Tr*9 microsatellite locus and TERG_6744 gene expression in *T. rubrum* genetic lineages. A. The number of repeats in STR*Tr*9 differs between ST1 and ST2 lineages. B. Relative expression of TERG_6744 gene, coding for fructose 1,6-bisphosphate aldolase, on a linear scale. It varies between strains irrespectively of their genetic lineage affiliation. There is no immediate consequence of STR*Tr*9 repeat number for TERG_06744 transcript level.

The analysis of molecular variance with 12 microsatellite loci revealed neglectable contribution of TERG_02941 793G subdivision to intergroup and overall variability (Table 3). With the removal of STR*Tr*9 from the analysis, the percent of variability, explained by the differences between genetic lineages, dropped drastically. But, this was accompanied by significant increase in explanatory power of TERG_02941 793G group subdivision. A half of the between-group variability in the dataset with 11 loci depended on it.

**Table 3.**
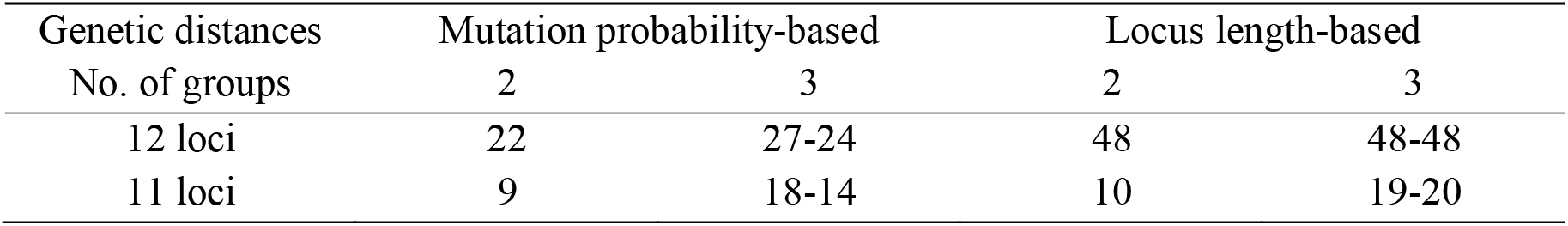
The analysis of molecular variance (AMOVA) in 70 *T. rubrum* isolates, performed on microsatellite polymorphism-based genetic distances. The numbers reflect percent of variability among TERG_02941 793A and 793G groups of isolates from the total amount of variability between and within groups. To obtain three groups, TERG_02941 793G isolates were further divided according to the topology of Bruvo distances-based network (left-hand numbers in columns with data on 3 lineages) or by Rozenfeld distances-based network (right-hand numbers).

In the sample of 70 isolates, the association index value 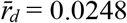 laid beyond the right boundary of the distribution of expected frequencies. The probability 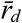 was estimated at 0.002, which made it possible to reject the null hypothesis of the absence of connections between the markers. Hence, the studied *T. rubrum* population was considered to be clonal. Also, six Russian isolates, subjected here to whole genome sequencing, harbored *MAT1-1* idiomorph of mating type locus. Simpson index was calculated on the basis of data from 11 neutral SSR loci. Its value within 95% confidence interval was estimated at 0.59-1.03 for ST1 lineage and 0.38-0.91 for ST2 lineage. The Fisher exact test showed no significant association between TERG_02941 genotypes and the localization of the lesion on patient’s body. Also, Yates-corrected *χ*^2^ test showed no connection between TERG_02941 genotype and a city of origin.

The CAAG repeats of STR*Tr*9 ended at the position −278 in the 5’-region of fructose 1,6-bisphosphate aldolase gene TERG_06744, wherein the adenine of the ATG translation initiation codon is considered +1. Therefore, we hypothesized that STR*Tr*9 locus promotes microevolution in *T. rubrum* through regulation of aldolase gene expression, which in its turn is one of the key enzymes of glycolysis.^42^ We determined relative TERG_06744 gene expression in six *T. rubrum* isolates. The isolate D15P44 with maximal length of STR*Tr*9 locus demonstrated the lowest level of TERG_06744 transcripts (Fig. 2B). The highest levels of TERG_06744 transcripts were observed in D15P62 isolate with the microsatellite length very close to minimal one. In the same time, three isolates with the same STR*Tr*9 length demonstrated significantly lower levels of TERG_06744 gene expression. Also, there were no clear differences between TERG_06744 transcript levels in isolates of different sequence types.

Russian sample of *T. rubrum* isolates was placed in global context by whole genome sequencing in six *T. rubrum* isolates. To get a representative sample, we selected isolates D15P33, D15P44, D15P50, D15P62, D15P71 and Y1013, covering the whole span of microsatellite distances-based Neighbour-Joining tree (not shown). These isolates were tested for susceptibility to antifungals (Table 4).

**Table 4.** Annotation and antifungal susceptibility data for six Russian *T. rubrum* isolates, subjected to whole genome sequencing.

| Isolate | Year of isolation | Place of isolation | Clinical picture | MIC<sub>50</sub> of antifungals (µg/ml) |  |  |  |  |
| --- | --- | --- | --- | --- | --- | --- | --- | --- |
|  |  |  |  | TER | ICZ | FCZ | POS | VRC |
| D15P33 | 2015 | St. Petersburg | OM, TP | ≤ 0.01 | 0.06 | 0.125 | 0.125 | 0.25 |
| D15P44 | 2015 | St. Petersburg | OM | ≤ 0.01 | 0.125 | 0.25 | 0.25 | 0.125 |
| D15P50 | 2016 | St. Petersburg | OM | ≤ 0.01 | 0.06 | 0.125 | 0.125 | 0.125 |
| D15P62 | 2016 | St. Petersburg | OM | ≤ 0.01 | 0.06 | 0.25 | 0.125 | 0.125 |
| D15P71 | 2016 | St. Petersburg | OM | ≤ 0.01 | 0.125 | 0.06 | 0.125 | 0.25 |
| Y1013 | 2017 | Yekaterinburg | OM | ≤ 0.01 | 0.125 | 0.25 | 0.25 | 0.125 |

Clinical picture: OM, toenail onychomycosis; TP, tinea pedis. Antifungals: TER, terbinafine; ICZ, itraconazole; FCZ, fluconazole; POS, posaconazole; VRC, voriconazole.

The phylogenomic tree included *T. rubrum sensu stricto* genomes from Europe, Asia and North America (Fig. 3) and contained well-supported monophyletic branch of ST1 isolates and basal paraphyletic ST2 branch. Both branches contained European and Asian isolates. In a representative selection of seven isolates from the study of Zheng et al.,^17^ cluster affiliation, determined in the original study, matched their TERG_02941/03298 genotypes. Depending on the presence of *T. violaceum* CMCC(F)T3l and *T. rubrum* “megninii” CBS 735.88 in the phylogenomic analysis, the depth of branches within *T. rubrum sensu stricto* clade related to *T. rubrum sensu stricto* — *T. soudanense* CBS 452.61 branch as 1:63 or 1:337 (with the two strains excluded).

**Figure 3.**
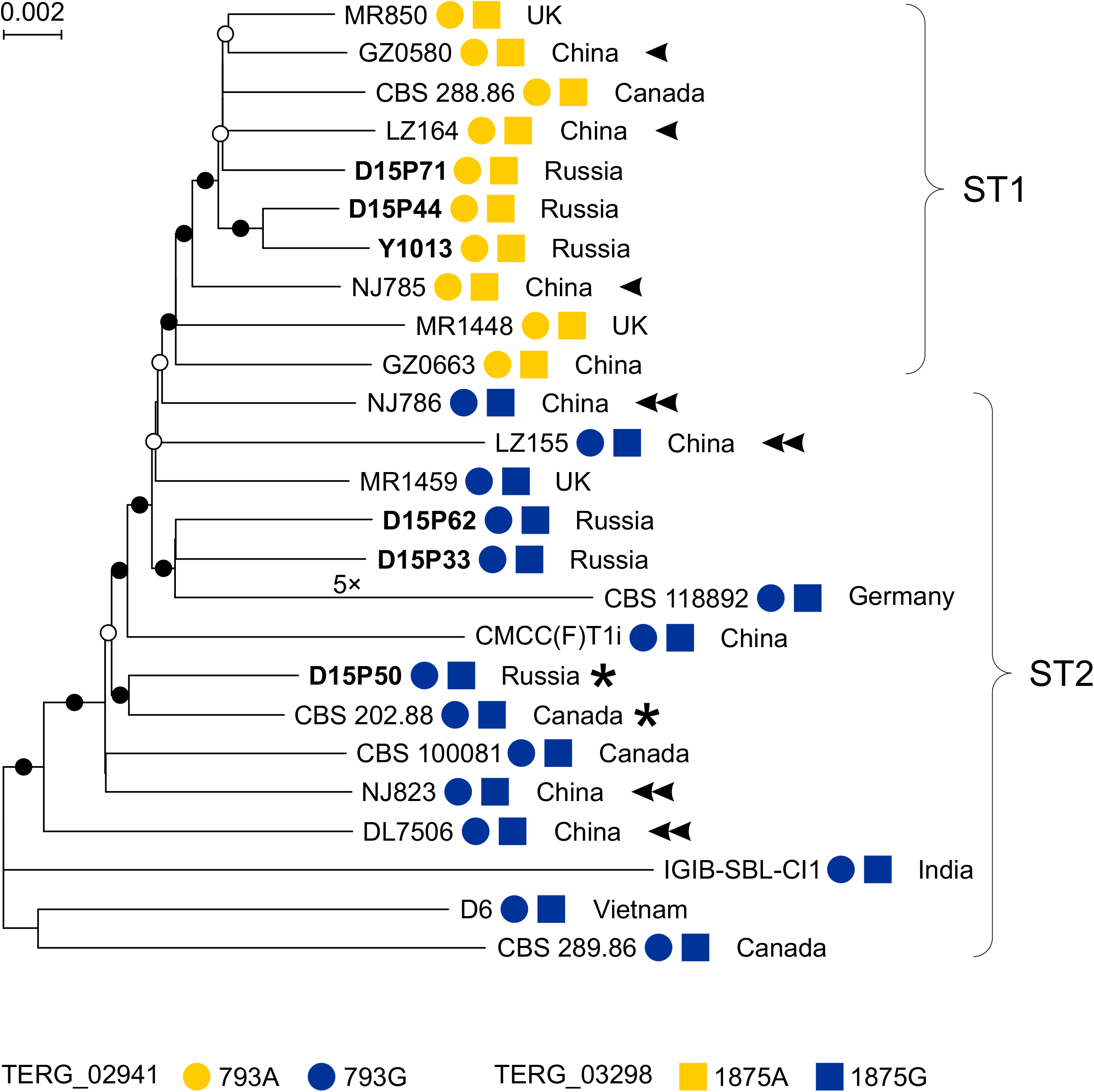
Phylogenomic tree of *T. rubrum* isolates. ST1 and ST2 lineages are defined by non-synonymous substitutions in TERG_02941 and TERG_03298 protein-coding loci. Single and double arrowheads mark Chinese isolates, fallen in two different clusters in the original publication by Zheng et al.^17^ The tree is rooted by *T. soudanense* CBS 452.61 whole genome sequence (the branch isn’t shown). Asterisks mark isolates with urease activity. Branch support values ≥ 75% are marked by open circles and values ≥ 95% are marked by solid circles. Newly sequenced isolates are given in bold. Russian population shares both lineages with West European and Asian countries.

Since urease activity has been regarded phylogenetically important trait in the dermatophytes,^43^ we assessed urease activity in our sample and found three urease-positive strains. For the isolate D15P50, whole genome sequence was available. There is an observation of the 68-bp intron and the 9-bp insertion in a urease-positive *T. rubrum* strain.^44^ However, in line with the results of Adamski et al.,^45^ we found identical urease gene sequences in all *T. rubrum sensu stricto* isolates, shown on Fig. 3, including urease-positive strains D15P50 and CBS 202.88. *Trichophyton rubrum* “megninii” CBS 735.88 harbored two point mutations in the urease gene, A685G in the intron and synonymous substitution G803A in the second exon. All our three isolates, positive for urease activity, came from the cases of toenail and foot skin infections and belonged to both major genetic lineages of *T. rubrum*.

## Discussion

Until recently, low genetic diversity of *T. rubrum* precluded persuasive description of its population structure. For this species, our earlier work seemingly represented the first successful attempt to obtain congruent results by two independent molecular strain typing techniques.^16^ However, we have not performed DNA sequencing of ribosomal ITS region to clarify whether the sample contained only *T. rubrum sensu stricto* isolates or not. Here we filled this gap by demonstrating that Russian population of the fungus is represented by only «classic» *T. rubrum* genotype. The two genetic lines of Russian *T. rubrum* population, found here, demonstrated significant divergence. The AMOVA analysis showed relatively high proportion of variability, explained by the division of *T. rubrum* sample into ST1 and ST2 groups, 22-48% depending on the distance metrics. It was not less, than in other studies on clonal filamentous ascomycetes with 19-21% of variability between groups^46,47^ or filamentous ascomycetes with mixed reproduction mode, with 2-13% of total variability explained by differences between intraspecific groups.^48,49^

Intraspecific genetic lines in fungi sometimes demonstrate restricted geographic ranges.^50–53^ Here, we compiled the map of global distribution of *T. rubrum* genetic lineages (Fig. 4). We largely built upon existing literature on NTS typing in *T. rubrum* and departed from the observation that single 450 bp PCR product of TRS-1 locus is associated with ST1 genetic lineage.^16^ We didn’t observe a specific distribution pattern of the two genetic lineages, but in most parts of the World, the derived ST1 lineage of *T. rubrum* presumably had higher prevalence and therefore demonstrated evolutionary success.

**Figure 4.**
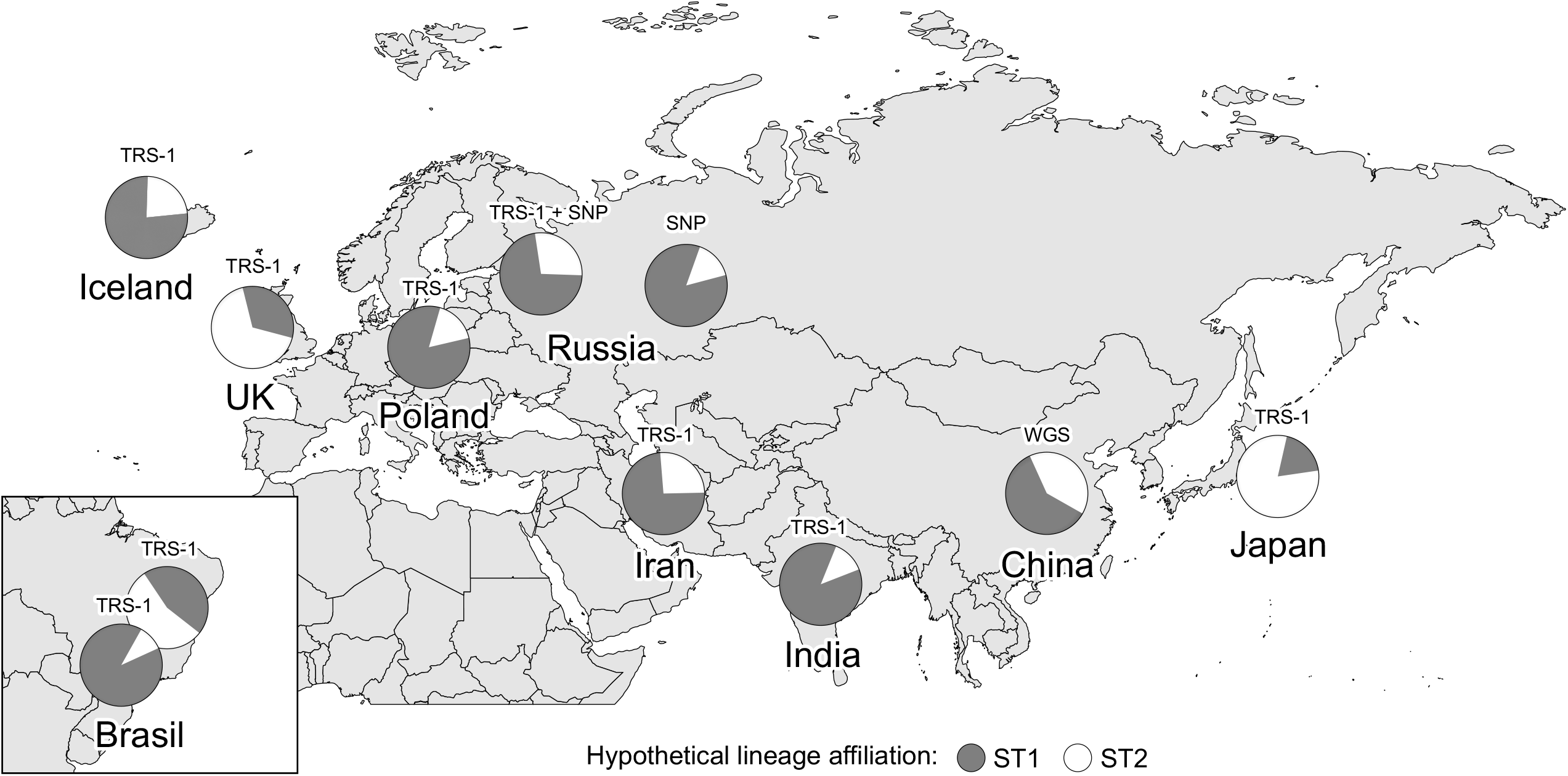
Hypothetical distribution of *Trichophyton rubrum* genetic lines. A total of 443 isolates, from 17 to 70 isolates for a country.^16,17,54–61^ The abbreviations above pie charts indicate the sources of variability: SNP, single nucleotide polymorphism of TERG_02941 and TERG_03298 genes; TRS-1, the length of TRS-1 locus of ribosomal non-transcribed spacer; WGS, whole genome sequences.

The affiliation of *T. rubrum* isolates to genetic lineages can be probed not only by NTS, SNP and WGS-based typing techniques, but also by our modified scheme of microsatellite analysis. But, this result may be of limited interest for healthcare practitioners, since there is seemingly no association of lineage affiliation and common isolate descriptors, like geography of origin and clinical picture of respective infection. Also, in the study of Zheng et al.,^17^ there was no association between cluster affiliation and antifungal susceptibility levels. It is widely accepted that truly independent species of clonal microbes occupy distinct ecological niches.^62–64^ In present study, there seemingly have been no clinical differences between the subpopulations, contrary to what we have claimed earlier.^16^ In this previous study, the association of STs and clinical forms of dermatophytosis might have occurred due to some random factors. From this perspective, the two genetic lineages of *T. rubrum sensu stricto* belong to the same fungal species.

The STR*Tr*9 locus was paramount for the analysis of the structure of the *T. rubrum* population by microsatellite analysis. Its length significantly differed between the isolates of the two genetic lines, implying the pressure of natural selection. There were significant differences in relative aldolase gene expression levels between different isolates. At the scale of our experiments, these differences could not be associated with population structure. Also, in our sample there were three urease-positive isolates, a proportion of 4% from the total. We did not find peculiarities in urease gene sequences of urease-positive isolates D15P50 and CBS 202.88. Positive strains belonged to both ST1 and ST2 groups. Therefore, our results suggest that urease-positive strains present in certain proportion in the population of the fungus, and represent natural variation of *T. rubrum sensu stricto*, not an independent entity. Lackner et al.^65^ obtained comparable results, studying virulence potential of closely related *Aspergillus* species from the section *Terrei*. They concluded that virulence is a strain-specific rather than species-specific feature. Lineage-specific differences in STR*Tr*9 locus length may still hallmark different adaptation strategies, given that glycolysis pathway regulation is connected with mechanisms of keratin degradation.^66^

Urease activity is an ancestral trait, commonly observable in geophilic dermatophytes^43^ and we expected to find it only in basal ST2 lineage. However, in our sample one of three urease-positive isolates belonged to ST1 lineage. Also, genetic diversity in the two lineages proved to be the same, which is contrary to what might be expected if ST2 is an ancestral lineage. It can be probably due to a sort of bottleneck, occurred in the population on the way from South-East Asia to Russia. In future studies, it is necessary to sample isolates from South and South-East Asia, in order to find the territory with highest genetic diversity and thereby conclusively determine the geographic region of the origin of the species.

Thus, we provided the description of the two intraspecific global lineages of *T. rubrum sensu stricto*. They can be revealed by SNP, SSR and WGS-based typing techniques (and NTS, with a certain amount of ambiguity). The species is polymorphic in terms of ecology and physiology, but this polymorphism is not associated with genetic population structure.

## Acknowledgements

The authors would like to thank the CBS-KNAW culture collection of Westerdijk Fungal Biodiversity Institute, Utrecht, the Netherlands, for providing the strain CBS 392.58. We also appreciate valuable comments from Adéla Čmoková and Ali Rezaei-Matehkolaei. This work was supported by Russian Foundation for Basic Research [grant number 18-34-00153].

## Author contributions

I.P. designed the study. I.P., Y.M., I.V., S.R. and T.B. performed the experiments. G.C. performed morphological species identification and maintained fungal cultures. I.P., D.A., V.Z. and Y.M. analyzed the data with the participation of A.T. N.V., A.T. and S.A. performed supervision and funding acquisition. I.P. wrote the manuscript with the help of V.Z.

## Data accessibility

Raw WGS reads were submitted to NCBI SRA database, BioProject PRJNA552357: the isolate D15P33, accession SRX6389058; D15P44, SRX6389060; D15P50, SRX6389057; D15P62, SRX6389059; D15P71, SRX6389062; Y1013, SRX6389061.

## Supplementary data

Supplement S1. Molecular strain typing in 70 *T. rubrum* isolates.

## References

1. Sigurgeirsson B, Baran R. The prevalence of onychomycosis in the global population: a literature study. J Eur Acad Dermatol Venereol. 2014;28(11):1480–1491. https://doi.org/10.1111/jdv.12323

2. Summerbell RC, Haugland RA, Li A, Gupta AK. rRNA gene internal transcribed spacer 1 and 2 sequences of asexual, anthropophilic dermatophytes related to *Trichophyton rubrum*. J Clin Microbiol. 1999;37(12):4005–4011.

3. Li HC, Bouchara JP, Hsu MM, Barton R, Su S, Chang TC. Identification of dermatophytes by sequence analysis of the rRNA gene internal transcribed spacer regions. J Med Microbiol. 2008;57(5):592–600. https://doi.org/10.1099/jmm.0.47607-0

4. Zhan P, Dukik K, Li D, et al. Phylogeny of dermatophytes with genomic character evaluation of clinically distinct *Trichophyton rubrum* and *T. violaceum*. Stud Mycol. 2018;89:153–175. https://doi.org/10.1016/j.simyco.2018.02.004

5. Su H, Packeu A, Ahmed SA, et al. Species distinction in the *Trichophyton rubrum* complex. J Clin Microbiol. 2019;57(9):e00352–19. https://doi.org/10.1128/jcm.00352-19

6. Packeu A, Stubbe D, Roesems S, et al. Lineages within the *Trichophyton rubrum* complex. Mycopathologia. 2020;185(1):123–136. https://doi.org/10.1007/s11046-019-00386-z

7. Gräser Y, Fröhlich J, Presber W, de Hoog S. Microsatellite markers reveal geographic population differentiation in *Trichophyton rubrum*. J Med Microbiol. 2007;56(8):1058–1065. https://doi.org/10.1099/jmm.0.47138-0

8. Gong J, Wu W, Ran M, et al. Population differentiation and genetic diversity of *Trichophyton rubrum* as revealed by highly discriminatory microsatellites. Fungal Genet Biol. 2016;95:24–29. https://doi.org/10.1016/j.fgb.2016.08.002

9. Gräser Y, Kühnisch J, Presber W. Molecular markers reveal exclusively clonal reproduction in *Trichophyton rubrum*. J Clin Microbiol. 1999;37(11):3713–3717.

10. Mirhendi H, Makimura K, de Hoog GS, et al. Translation elongation factor 1-α gene as a potential taxonomic and identification marker in dermatophytes. Med Mycol. 2015;53(3):215–224. https://doi.org/10.1093/mmy/myu088

11. Pchelin IM, Zlatogursky VV, Rudneva MV, et al. Reconstruction of phylogenetic relationships in dermatomycete genus *Trichophyton* Malmsten 1848 based on ribosomal internal transcribed spacer region, partial 28S rRNA and beta-tubulin genes sequences. Mycoses. 2016;59(9):566–575. https://doi.org/10.1111/myc.12505

12. Li W, Metin B, White TC, Heitman J. Organization and evolutionary trajectory of the mating type (MAT) locus in dermatophyte and dimorphic fungal pathogens. Eukaryot Cell. 2010;9(1):46–58. https://doi.org/10.1128/EC.00259-09

13. Kano R, Isizuka M, Hiruma M, Mochizuki T, Kamata H, Hasegawa A. Mating type gene (*MAT1-1*) in Japanese isolates of *Trichophyton rubrum*. Mycopathologia. 2013;175(1-2):171–173. https://doi.org/10.1007/s11046-012-9603-2

14. Persinoti GF, Martinez DA, Li W, et al. Whole-genome analysis illustrates global clonal population structure of the ubiquitous dermatophyte pathogen *Trichophyton rubrum*. Genetics. 2018;208(4):1657–1669. https://doi.org/10.1534/genetics.117.300573

15. Arabatzis M, Velegraki A, Kantardjiev T, Stavrakieva V, Rigopoulos D, Katsambas A. First report on autochthonous urease-positive *Trichophyton rubrum* (*T. raubitschekii*) from South-east Europe. Br J Dermatol. 2005;153(1):178–182. https://doi.org/10.1111/j.1365-2133.2005.06615.x

16. Pchelin IM, Azarov DV, Chilina GA, Dmitriev KA, Vasilyeva NV, Taraskina AE. Single-nucleotide polymorphism in a local population of *Trichophyton rubrum*. Med Mycol. 2018;56(1):125–128. https://doi.org/10.1093/mmy/myx009

17. Zheng H, Blechert O, Mei H, et al. Whole-genome resequencing of *Trichophyton rubrum* provides insights into population differentiation and drug resistance. Mycopathologia. 2020;185(1):103–112. https://doi.org/10.1007/s11046-019-00384-1

18. Martinez DA, Oliver BG, Gräser Y, et al. Comparative genome analysis of *Trichophyton rubrum* and related dermatophytes reveals candidate genes involved in infection. MBio. 2012;3(5):e00259–12. https://doi.org/10.1128/mBio.00259-12

19. Latka C, Dey SS, Mahajan S, et al. Genome sequence of a clinical isolate of dermatophyte, *Trichophyton rubrum* from India. FEMS Microbiol Lett. 2015;362(8):fnv039. https://doi.org/10.1093/femsle/fnv039

20. White TJ, Bruns TD, Lee SB, Taylor JW. Amplification and Sequencing of Fungal Ribosomal RNA Genes for Phylogenetics, In PCR Protocols and Applications: A Laboratory Manual. New York, NY: Academic Press; 1990:315–322.

21. Jacob TR, Peres NT, Persinoti GF, et al. *rpb2* is a reliable reference gene for quantitative gene expression analysis in the dermatophyte *Trichophyton rubrum*. Med Mycol. 2012;50(4):368–377. https://doi.org/10.3109/13693786.2011.616230

22. Arendrup MC, Meletiadis J, Mouton JW, et al. EUCAST definitive document E.Def.3.1 method for the determination of broth dilution minimum inhibitory concentrations of antifungal agents for conidia forming moulds. EUCAST. 2017.

23. Ye J, Coulouris G, Zaretskaya I, Cutcutache I, Rozen S, Madden T. Primer-BLAST: A tool to design target-specific primers for polymerase chain reaction. BMC Bioinformatics. 2012;13:134. https://doi.org/10.1186/1471-2105-13-134

24. Pchelin IM, Azarov DV, Churina MA, et al. Whole genome sequence of first *Candida auris* strain, isolated in Russia. Med Mycol. 2020;58(3):414–416. https://doi.org/10.1093/mmy/myz078

25. Andrews S. FastQC: a quality control tool for high throughput sequence data. 2010. http://www.bioinformatics.babraham.ac.uk/projects/fastqc. Accessed: 22.02.2020.

26. Bolger AM, Lohse M, Usadel B. Trimmomatic: a flexible trimmer for Illumina sequence data. Bioinformatics. 2014;30(15):2114–2120. https://doi.org/10.1093/bioinformatics/btu170

27. Nurk S, Bankevich A, Antipov D, et al. Assembling genomes and mini-metagenomes from highly chimeric reads. In M. Deng, R. Jiang, F. Sun, & X. Zhang (Eds.), Annual International Conference on Research in Computational Molecular Biology (pp. 158–170). Berlin, Heidelberg: Springer; 2013. https://doi.org/10.1007/978-3-642-37195-0_13

28. Treangen TJ, Ondov BD, Koren S, Phillippy AM. The Harvest suite for rapid core-genome alignment and visualization of thousands of intraspecific microbial genomes. Genome Biol. 2014;15(11):524. https://doi.org/10.1186/s13059-014-0524-x

29. Price MN, Dehal PS, Arkin AP. FastTree 2–approximately maximum-likelihood trees for large alignments. PLoS One. 2010;5:e9490. https://doi.org/10.1371/journal.pone.0009490

30. Bruvo R, Michiels NK, D’Souza TG, Schulenburg H. A simple method for the calculation of microsatellite genotype distances irrespective of ploidy level. Mol Ecol. 2004;13(7):2101–2106. https://doi.org/10.1111/j.1365-294X.2004.02209.x

31. Clark L, Jasieniuk M. Polysat: an R package for polyploid microsatellite analysis. Mol Ecol Resour. 2011;11(3):562–566. https://doi.org/10.1111/j.1755-0998.2011.02985.x

32. Rozenfeld AF, Arnaud-Haond S, Hernández-García E, et al. Spectrum of genetic diversity and networks of clonal organisms. J R Soc Interface. 2007;4(17):1093–1102. https://doi.org/10.1098/rsif.2007.0230

33. Huson DH, Bryant D. Application of phylogenetic networks in evolutionary studies. Mol Biol Evol. 2006;23(2):254–267. https://doi.org/10.1093/molbev/msj030

34. Tamura K, Stecher G, Peterson D, Filipski A, Kumar S. MEGA6: molecular evolutionary genetics analysis version 6.0. Mol Biol Evol. 2013;30(12):2725–2729. https://doi.org/10.1093/molbev/mst197

35. Kamvar ZN, Tabima JF, Grünwald NJ. Poppr: an R package for genetic analysis of populations with clonal, partially clonal, and/or sexual reproduction. PeerJ. 2014;2:e281. https://dx.doi.org/10.7717/peerj.281

36. Peakall R, Smouse PE. GENALEX 6: genetic analysis in Excel. Population genetic software for teaching and research. Mol Ecol Notes. 2006;6(1):288–295. https://doi.org/10.1111/j.1471-8286.2005.01155.x

37. Peakall R, Smouse PE. GenAlEx 6.5: genetic analysis in Excel. Population genetic software for teaching and research – an update. Bioinformatics. 2012;28(19):2537–2539. https://doi.org/10.1093/bioinformatics/bts460

38. Zhang Z, Schwartz S, Wagner L, Miller W. A greedy algorithm for aligning DNA sequences. J Comput Biol. 2000;7(1-2):203–214. https://doi.org/10.1089/10665270050081478

39. Kano R, Kawasaki M, Mochizuki T, Hiruma M, Hasegawa A. Mating genes of the *Trichophyton mentagrophytes* complex. Mycopathologia. 2012;173(2-3):103–112. https://doi.org/10.1007/s11046-011-9487-6

40. Pchelin IM, Kryuchkova MA, Bogdanova TV, et al. We need more powerful microsatellite assay for population genetic studies of *Trichophyton rubrum*. Med Mycol. 2018;56(S2):S119.

41. Leliaert F, Verbruggen H, Vanormelingen P, et al. DNA-based species delimitation in algae. Eur J Phycol. 2014;49(2):179–196. https://dx.doi.org/10.1080/09670262.2014.904524

42. Nelson DL, Cox MM. Lehninger principles of biochemistry. New York: Freeman; 2008.

43. Summerbell RC. Form and function in the evolution of dermatophytes. In: Kushwaha RKS, Guarro J, editors. Biology of dermatophytes and other keratinophilic fungi. Rev Iberoam Micol. 2000;17(Suppl):30–43.

44. Hiruma M, Kano R, Sugita T, Mochizuki T, Hasegawa A, Hiruma M. Urease gene of *Trichophyton rubrum* var. *raubitschekii*. J Dermatol. 2013;40:111–113. https://doi.org/10.1111/1346-8138.12017

45. Adamski Z, Kowalczyk MJ, Adamska K, et al. The first non-African case of *Trichophyton rubrum* var. *raubitschekii* or a urease-positive *Trichophyton rubrum* in Central Europe? Mycopathologia. 2014;178(1-2):91–96. https://doi.org/10.1007/s11046-014-9751-7

46. Ahmadpour A, Castell-Miller C, Javan-Nikkhah M, et al. Population structure, genetic diversity, and sexual state of the rice brown spot pathogen *Bipolaris oryzae* from three Asian countries. Plant Pathol. 2018;67(1):181–192. https://doi.org/10.1111/ppa.12714

47. Singh N, Anand G, Kapoor R. Virulence and genetic diversity among *Fusarium oxysporum* f. sp. *carthami* isolates of India using multilocus RAPD and ISSR markers. Trop Plant Pathol. 2019;44(5):409–422. https://doi.org/10.1007/s40858-019-00303-1

48. Sexton AC, Howlett BJ. Microsatellite markers reveal genetic differentiation among populations of *Sclerotinia sclerotiorum* from Australian canola fields. Curr Genet. 2004;46(6):357–365. https://doi.org/10.1007/s00294-004-0543-3

49. Kashyap PL, Kumar S, Kumar RS, et al. Identification of novel microsatellite markers to assess the population structure and genetic differentiation of *Ustilago hordei* causing covered smut of barley. Front Microbiol. 2020;10:2929. https://doi.org/10.3389/fmicb.2019.02929

50. Abdel-Rahman SM, Sugita T, González GM, et al. Divergence among an international population of *Trichophyton tonsurans* isolates. Mycopathologia. 2010;169(1):1–13. https://doi.org/10.1007/s11046-009-9223-7

51. Mironenko NV, Baranova OA, Kovalenko NM, Mikhailova LA, Rosseva LP. Genetic structure of the Russian populations of *Pyrenophora tritici-repentis*, determined by using microsatellite markers. Russ J Genet. 2016;52(8):771–779. https://doi.org/10.1134/S1022795416080093

52. Gultyaeva EI, Aristova MK, Shaidayuk EL, et al. Genetic differentiation of *Puccinia triticina* Erikss. in Russia. Russ J Genet. 2017;53(9):998–1005. https://doi.org/10.1134/S1022795417070031

53. Taghipour S, Pchelin IM, Zarei Mahmoudabadi A, et al. *Trichophyton mentagrophytes* and *T. interdigitale* genotypes are associated with particular geographic areas and clinical manifestations. Mycoses. 2019;62(11):1084–1091. https://doi.org/10.1111/myc.12993

54. Jackson CJ, Barton RC, Kelly SL, Evans EG. Strain identification of *Trichophyton rubrum* by specific amplification of subrepeat elements in the ribosomal DNA nontranscribed spacer. J Clin Microbiol. 2000;38(12):4527–4534.

55. Baeza LC, Matsumoto MT, Almeida AMF, Mendes-Giannini MJS. Strain differentiation of *Trichophyton rubrum* by randomly amplified polymorphic DNA and analysis of rDNA nontranscribed spacer. J Med Microbiol. 2006;55(Pt 4):429–436. doi:10.1099/jmm.0.46236-0

56. de Assis Santos D, de Carvalho Araújo RA, Kohler LM, Machado-Pinto J, Hamdan JS, Cisalpino PS. Molecular typing and antifungal susceptibility of *Trichophyton rubrum* isolates from patients with onychomycosis pre- and post-treatment. Int J Antimicrob Agents. 2007;29(5):563–569. doi:10.1016/j.ijantimicag.2006.09.028

57. Rad MM, Mohammadi AMA, Barton RC. PCR typing of *Trichophyton rubrum* isolates by specific amplification of subrepeat elements in ribosomal DNA nontranscribed spacer. Iranian J Dermatol. 2008;11(1):17–20.

58. Hryncewicz-Gwóźdź A, Jagielski T, Sadakierska-Chudy A, et al. Molecular typing of *Trichophyton rubrum* clinical isolates from Poland. Mycoses. 2011;54(6):e726–e736. doi:10.1111/j.1439-0507.2010.02007.x

59. Takahashi I, Fukushima K, Miyaji M, Nishimura K, Asano K, Iizuka H. Species identification and strain typing of dermatophytes by single-strand conformation polymorphism (SSCP) analysis of the ribosomal DNA and polymerase chain reaction analysis of subrepeat elements in the intergenic spacer region of *Trichophyton rubrum*. Asahikawa Medical University Repository. 2015;15:27–36.

60. Ramaraj V, Vijayaraman RS, Elavarashi E, Rangarajan S, Kindo AJ. Molecular strain typing of clinical isolates, *Trichophyton rubrum* using non transcribed spacer (NTS) region as a molecular marker. J Clin Diagn Res. 2017;11(5):DC04–DC09. doi:10.7860/JCDR/2017/21994.9843

61. Suzuki S, Mano Y, Furuya N, Fujitani K. Molecular epidemiological analysis of the spreading conditions of *Trichophyton* in long-term care facilities in Japan. Jpn J Infect Dis. 2018;71(6):462–466. doi:10.7883/yoken.JJID.2018.090

62. Tibayrenc M. The species concept in parasites and other pathogens: a pragmatic approach? Trends Parasitol. 2006;22(2):66–70. https://doi.org/10.1016/j.pt.2005.12.010

63. Achtman M, Wagner M. Microbial diversity and the genetic nature of microbial species. Nat Rev Microbiol. 2008;6(6):431–440. https://doi.org/10.1038/nrmicro1872

64. Giraud T, Refrégier G, Le Gac M, de Vienne DM, Hood ME. Speciation in fungi. Fungal Genet Biol. 2008;45(6):791–802. https://doi.org/10.1016/j.fgb.2008.02.001

65. Lackner M, Obermair J, Naschberger V, et al. Cryptic species of *Aspergillus* section *Terrei* display essential physiological features to cause infection and are similar in their virulence potential in *Galleria mellonella*. Virulence. 2019;10(1):542–554. https://doi.org/10.1080/21505594.2019.1614382

66. Martins MP, Rossi A, Sanches PR, Bortolossi JC, Martinez-Rossi NM. Comprehensive analysis of the dermatophyte *Trichophyton rubrum* transcriptional profile reveals dynamic metabolic modulation. Biochem J. 2020;477(5):873–885. doi:10.1042/BCJ20190868

